# Supra second timing reflects oscillatory and aperiodic EEG dynamics

**DOI:** 10.64898/2026.07.02.736244

**Authors:** Lorenzo Guarnieri, Ayelet N. Landau

**Affiliations:** Hebrew University of Jerusalem (HUJI); University College London (UCL)

## Abstract

The perception of timing has been investigated across short and long timescales, with the latter usually underrepresented. Previous accounts of supra-second timing have often emphasised dedicated neural signals, yet whether timing performance over longer durations reflects specialised temporal mechanisms or domain-general neural excitability remains unclear. We examined this question in an interval reproduction task (2–4 s) using EEG, decomposing neural activity into oscillatory (alpha, theta) and aperiodic components and relating both to behavioural performance. At the neural level, separate analyses were performed on the interval encoding epoch and the following delay period. Time-resolved analyses revealed a coordinated decrease in posterior alpha power and aperiodic offset during interval encoding. In contrast, interval duration did not produce consistent modulation of oscillatory or aperiodic activity during the delay period, providing limited support for a workload-based account of time duration. Across participants, higher baseline alpha power and aperiodic offset were associated with better timing accuracy, whereas trial-by-trial fluctuations in aperiodic activity, and to a lesser extent alpha power, predicted single-trial reproductions. The results suggest that temporal behaviour in the supra-second range is shaped by domain-general neural activity sustaining goal-directed task engagement— with oscillatory and aperiodic dynamics serving as complementary indices of this broader excitability state.

## Introduction

A longstanding question in neuroscience concerns the mechanisms through which the brain represents time across different timescales (Buhusi & Meck, 2005; Buzsáki, 2025). Classic accounts have often distinguished sub-second timing, which is thought to rely more strongly on sensory and automatic processes, from supra-second timing, which places greater demands on attention, working memory, and goal maintenance (Lewis & Miall, 2003). Studying supra-second timing presents unique challenges: behavioural and neural noise increase with interval duration (Duffy et al., 2025; Malapani & Fairhurst, 2002), and the sustained engagement over longer intervals makes it difficult to isolate timing mechanisms from the broader cognitive processes that necessarily accompany them (Buhusi & Meck, 2005). Rather than treating this entanglement as a confound, we take it as a starting point. We propose that supra-second timing is better understood not as the output of a dedicated temporal mechanism, (e.g., the post-interval P200/300; Damsma et al., 2021; Kononowicz & van Rijn, 2014), but as an emergent consequence of the neural activity supporting attention, working memory, and goal maintenance over extended periods. Theoretical models further suggest that cognitive processes, such as attention and memory, fluctuate dynamically over short and long timescales, alternating between high and low efficacy (Corriveau et al., 2025; Esterman & Rothlein, 2019; Fiebelkorn & Kastner, 2019; Herweg et al., 2020; Landau, 2018; Madore et al., 2022; Re et al., 2025; Rosenberg, 2025), and that this structured process may disperse or change during prolonged task engagement (Hemmerich, Luna, et al., 2025; Sharpe & Tyndall, 2025). We therefore examine supra-second timing performance through the lens of neural states that govern sustained cognitive engagement during temporal encoding. Focusing on the neural level, several candidate mechanisms may play a key role in processes unfolding over longer timescales. Alpha-band (8-12 Hz) activity has been associated with fluctuations in attentional state, tonic alertness, and the regulation of cortical excitability, whereas frontal theta (4-7 Hz) has been linked to cognitive control, working memory, and task demands (Cavanagh & Frank, 2014; Chikhi et al., 2022; Herweg et al., 2020; Klimesch, 2012; Sadaghiani & Kleinschmidt, 2016). Moreover, oscillatory accounts have provided an important framework for understanding temporal cognition from multiple perspectives (Damsma et al., 2021; Kononowicz et al., 2018; McCone et al., 2024; Wiener et al., 2018). In addition, recent work has highlighted the cognitive role of the aperiodic component (or 1/f) of the EEG power spectrum (Donoghue et al., 2020; Hemmerich, Santoni, et al., 2025). Rather than reflecting background noise, aperiodic activity has been shown to vary systematically with perception (Cunningham et al., 2023; Deodato & Melcher, 2024; Gyurkovics et al., 2022; Kałamała et al., 2024), working memory (Bender et al., 2025; Engen et al., 2026; Frelih et al., 2025; Preston et al., 2025; Virtue-Griffiths et al., 2025), and in some cases to account for effects previously attributed exclusively to oscillatory changes (Engen et al., 2026; Herzog et al., 2024; Pei et al., 2023; Zhang et al., 2023). This has motivated a broader view in which oscillatory and aperiodic activity provide complementary information about neural population dynamics and behaviour (although note Karvat et al., 2026). This complementary framework has not previously been applied to timing research, though we hypothesize that it might play a key role in sustaining neural excitability during long interval encoding. Accordingly, we tested whether supra-second timing performance reflects the state of ongoing neural activity supporting long-term task engagement, by jointly analysing oscillatory and aperiodic EEG dynamics during a reproduction task. We addressed this across two levels of analysis — individual differences in brain– behaviour relationships and moment-to-moment trial-by-trial fluctuations — through three specific questions. First, we asked whether oscillatory power (alpha and theta) and aperiodic activity scale with the encoding of different interval durations. Second, we inquired whether these neural dynamics scale with interval duration during a delay period, while different durations are maintained for subsequent reproduction. Third, at the level of individual differences and trial-by-trial variability, we asked how individual neural state and moment-to-moment fluctuations in oscillatory and aperiodic activity predict timing behaviour. Together, these questions suggest that supra-second timing as an emergent property of the neural states that regulate sustained cognitive engagement and resource allocation over longer periods.

## Results

### Behavioural results

Participants performed a temporal reproduction task. Empty intervals were presented using brief stimuli (i.e., markers) lasting 2, 3 or 4 seconds. Experimental design and behavioural results are depicted in Fig. 1 and further detailed in the methods section. To characterise individual timing behaviour, we fitted a Bayesian observer model (Cicchini et al., 2012; Jazayeri & Shadlen, 2010) to the reproduction data (see Methods). The group-level prior mean was estimated at 2.41 s (95% CI [1.96, 2.83]), indicating a general tendency to under-reproduce time. The group-level mean Weber constant was 1.02 (95% CI [0.21, 3.07]), consistent with scalar variability, whereby temporal noise increases proportionally with interval duration (Malapani & Fairhurst, 2002). Substantial between-subject variability was observed in the inferred prior mean (SD = 1.18, 95% CI [0.87, 1.6]), suggesting marked individual differences in prior-driven biases. To capture these individual differences more directly, we derived two complementary summary metrics from the model and behavioural data: (i) central tendency, indexing the relative influence of prior expectations on temporal estimates (Supp. Fig. 1A), and (ii) the behavioural slope, reflecting the gain with which encoded intervals are translated into reproduced durations (Supp. Fig. 1B). These measures dissociate bias (prior-driven compression toward the mean) from scaling (sensitivity to interval magnitude) and therefore provide a compact description of individual timing strategies.

**Figure 1:**
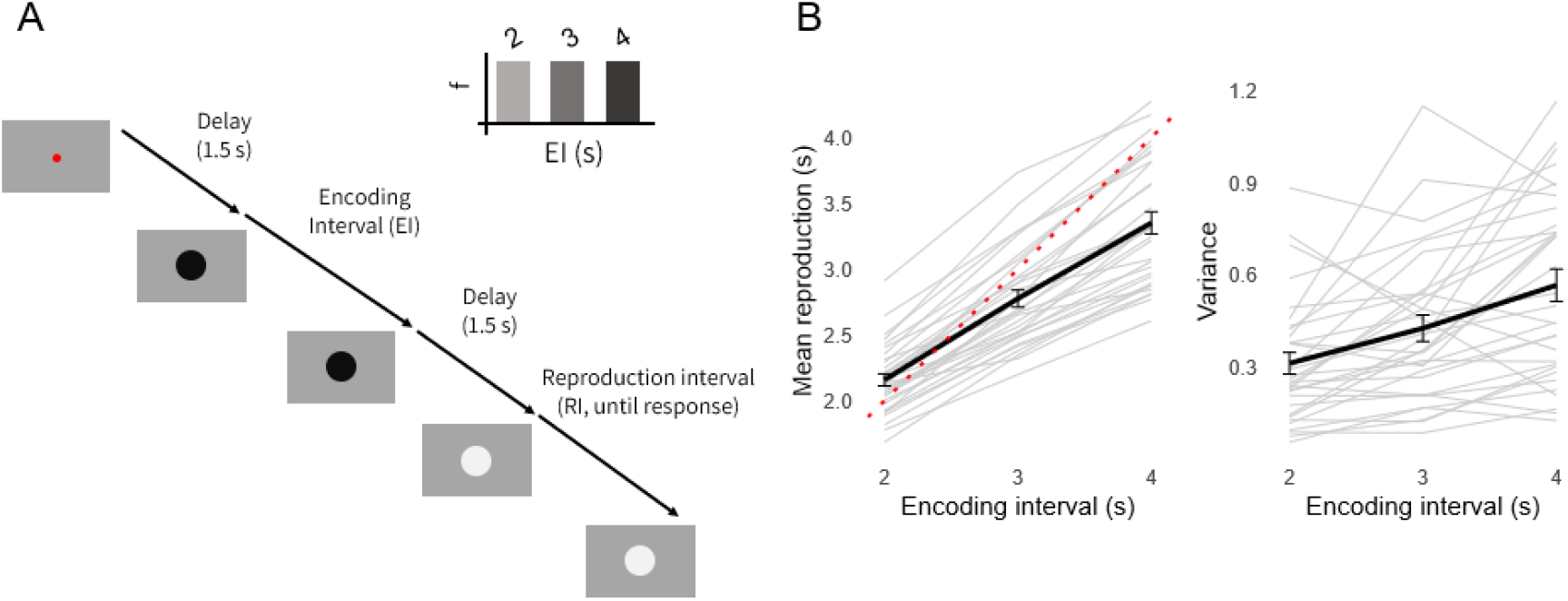
experimental design and behavioural results. (A) Schematic of the time reproduction task. Participants encoded one of three possible time intervals from the prior uniform distribution (2, 3, or 4 seconds), followed by a fixed delay of 1.5 seconds. They then reproduced the encoded interval by pressing a response key. B) Mean reproduction and variance as function of the encoded interval. The red dashed line represents the identity line (veridical performance). In both plots, grey lines indicate individual participants, while the solid black line shows the group average (± 1 SEM).

### Oscillatory and aperiodic activity modulation during the interval encoding and maintenance

To characterise the temporal dynamics of oscillatory and aperiodic activity during task-related phases, we performed a time-resolved spectral parameterization (Donoghue et al., 2020) using a sliding-windows approach (Bender et al., 2025). For each subject, channel, and interval condition, EEG data were analysed across the full encoding and delay period of the three interval conditions (−0.5 to 3.5 s, −0.5 to 4.5 s, and −0.5 to 5.5 s relative to EI onset), using overlapping windows of 0.5 s shifted in steps of 0.1 s. Within each window, FOOOF parameters were estimated, including oscillatory (alpha and theta) activity as well as aperiodic offset and exponent (refer to Methods for the parameter specifications). Resulting time series and topographies are shown in Fig. 2. To assess oscillatory and aperiodic modulation by interval duration, regression analyses with permutation were performed over the final second of each encoding interval (EI) and over the entire delay period (1.5 s). During the EI, alpha power significantly decreased as a function of interval duration (t = −3.945, p < 0.001), indicating reduced posterior alpha activity for longer intervals. This effect was not observed during the delay period (p > 0.1). In contrast, theta activity did not show significant modulation with interval duration during interval encoding (p > 0.1), nor did it exhibit robust effects during the delay, a trend in frontal channels that did not survive correction for multiple comparisons. Analysis of aperiodic activity revealed that offset significantly decreased as a function of encoding interval in occipital channels (t = −2.516, p = 0.0147), whereas the aperiodic exponent did not show significant modulation during encoding (p > 0.1).

**Figure 2:**
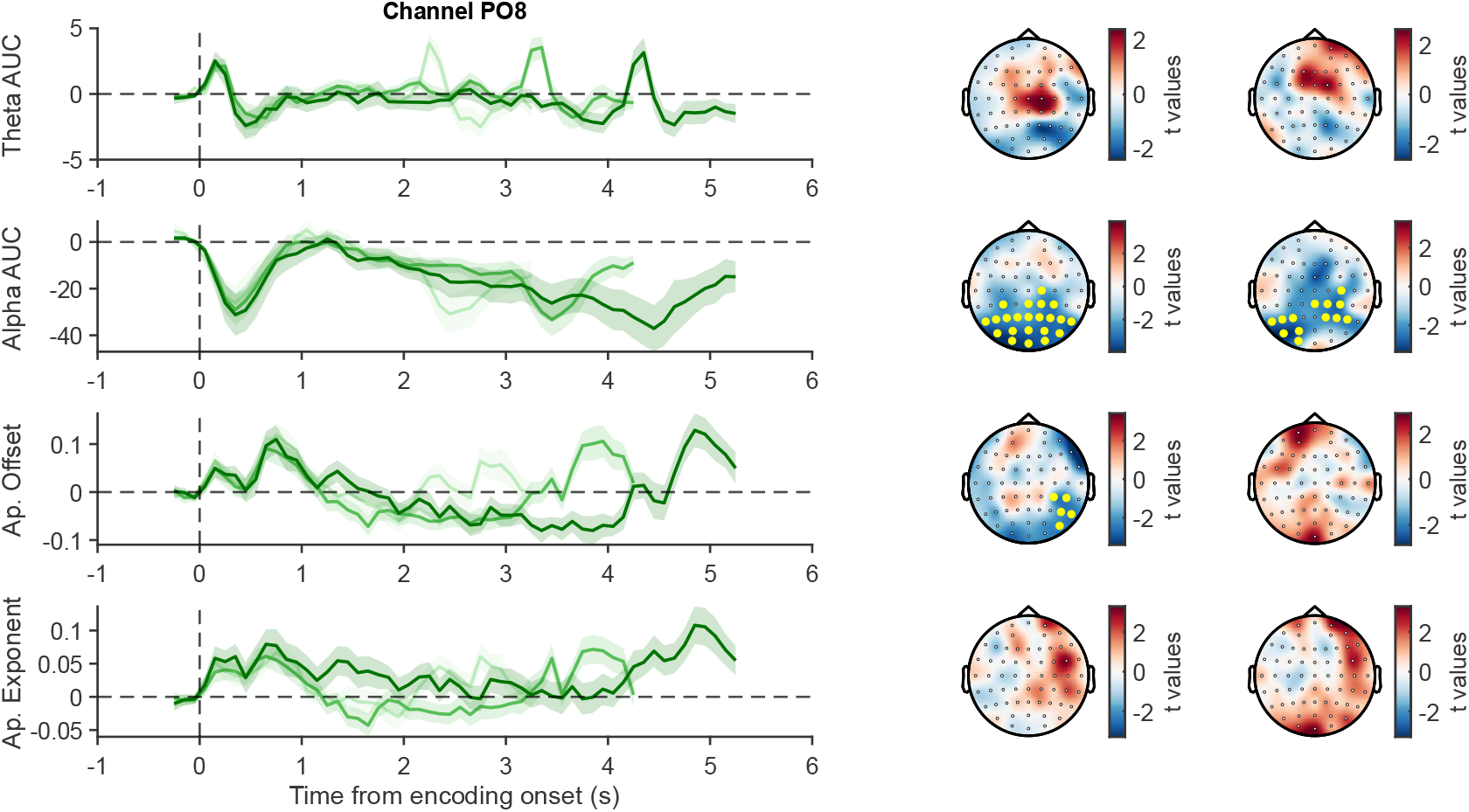
Temporal dynamics of oscillatory and aperiodic EEG components during encoding and delay. Left panels: Time-resolved changes at a posterior electrode (PO8) aligned to encoding onset (0 s). From top to bottom, rows correspond to: Theta AUC, Alpha AUC, aperiodic offset, and aperiodic exponent. Lines indicate the grand average (± 1 SEM), with darker traces indicating longer intervals. Data are baselined using a -0.5-0 s window before EI onset. Right panels: Scalp topographies of *t*-values for the final 1 s window of the encoding interval (EI) and the 1.5 s delay periods. The colour in the topographies denotes t-values, calculated as the regression estimates (across the intervals) divided by their standard error (β/SE(β)). Filled yellow markers denote significant channels (p < 0.05, cluster corrected).

### Individual differences in baseline alpha power and aperiodic offset predict timing accuracy

Previous work has emphasized the importance of understanding how the relationship between neural activity and behavioural performance varies across individuals (Deodato & Melcher, 2024; Glicksohn, 2024; Haegens et al., 2014; Madore et al., 2020). In this context, neural activity at task baseline or during rest can provide a useful index of ongoing brain state that is not directly tied to specific task events. Here, we examined whether baseline EEG activity, measured prior to the encoding interval onset, could account for individual differences in timing performance. For each subject, oscillatory (alpha, theta) and aperiodic (offset, exponent) parameters were extracted from a 1-s pre-stimulus baseline window preceding encoding interval onset. Notably, oscillatory alpha peaks were identifiable in nearly all participants (31/32), whereas clear frontal theta peaks were observed in less than half of the sample (11/32), consistent with recent reports highlighting substantial inter-individual variability and limited detectability of frontal theta oscillations (Engen et al., 2026). These baseline measures were then correlated with the individual behavioural parameters, namely individual slope (indexing timing accuracy) and central tendency (see Behavioural Results). Baseline alpha power showed a robust positive correlation with individual slope (r = 0.516, p = 0.0058), indicating that higher posterior alpha activity was associated with more veridical timing performance. In contrast, theta activity did not show a significant relationship with slope (r = 0.077, p = 0.5826). Furthermore, neither aperiodic offset (r = 0.183, p = 0.346) nor the exponent (r = -0.059, p = 0.5897) show any significant correlation with the Slope. Full correlation matrices and topographies are reported in Fig. 3. For what concerned the relation between the spectral measures and the central tendency, neither alpha (r = -0.189, p = 0.3456) nor theta (r = -0.092, p = 0.5362) showed significant effects. Aperiodic exponent was also not associated with central tendency (r = -0.12, p = 0.4149), while the offset showed a small significant cluster in parietal channels (r =-0.55, p = 0.0013). Full correlation matrices and topographies are reported in Supp. Fig. 2.

**Figure 3:**
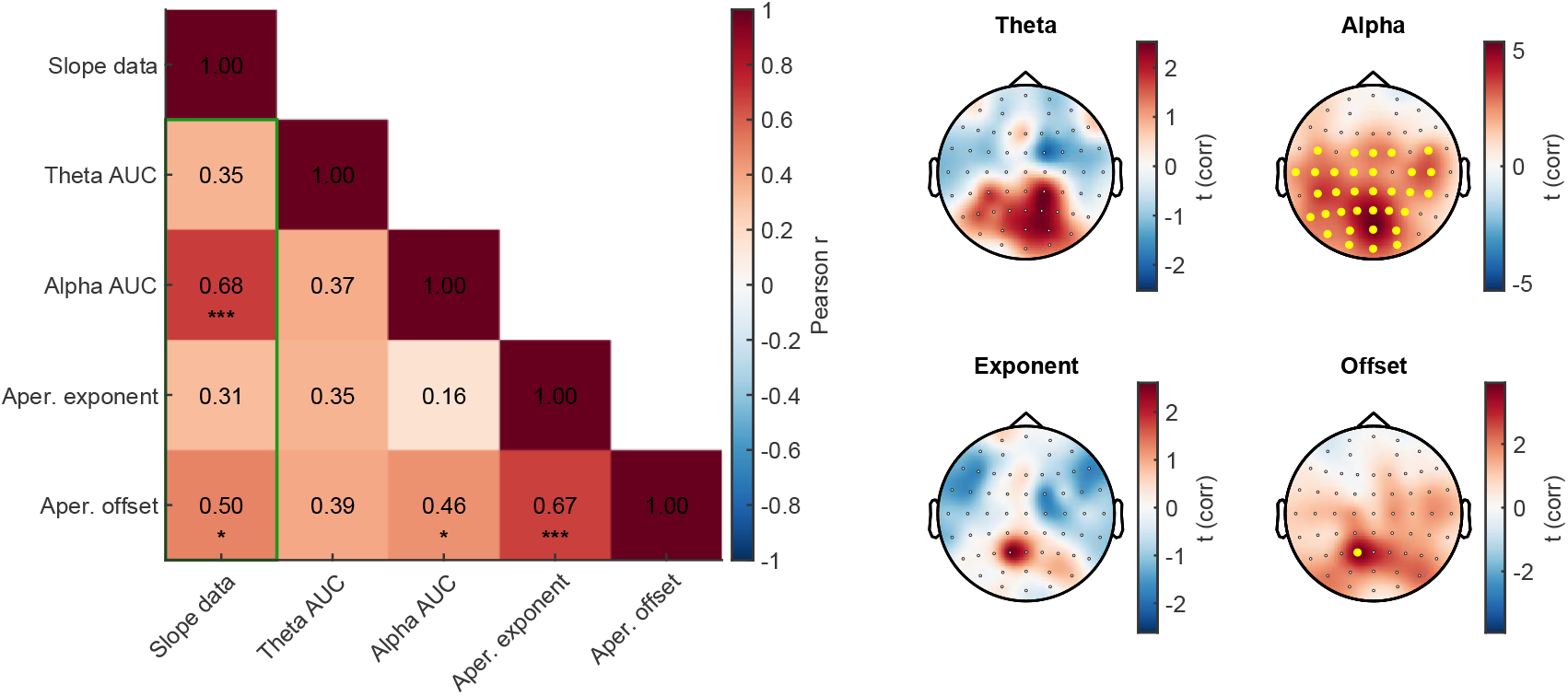
Relationship between behavioural timing metrics and EEG spectral components. Left panel: Pearson correlation matrix across subjects (N = 32) including behavioural slope (Slope_data), theta power (Theta AUC), alpha power (Alpha AUC), aperiodic exponent, and aperiodic offset, averaged across a posterior electrode cluster (CPz, P1, Pz, P2, POz). Green-outlined correlations are represented in the topographies on the right. Significant correlations are denoted by asterisks (* p < 0.05, ** p < 0.01, *** p < 0.001; FDR corrected). Right panels: Scalp topographies of channel-wise correlation t-values between behavioural slope (Slope_data) and EEG metrics. Top row: correlations with Theta AUC (left) and Alpha AUC (right). Bottom row: correlations with aperiodic exponent (left) and aperiodic offset (right). Yellow markers denote channels reaching statistical significance.

### Higher oscillatory alpha power and aperiodic activity predict longer single-trial reproductions

To assess the trial-by-trial contribution of neural activity to behavioural output, we conducted a linear mixed-effects analysis (see Methods) relating single-trial oscillatory and aperiodic parameters to reproduction times across the full duration of each encoding interval (2, 3, and 4 s). The full encoding interval was included as a proxy of the cognitive and neural dynamics evolving over time, which we hypothesised may ultimately shape the final reproduction. The spatial distribution of significant effects is shown in Fig. 4. At the group level, alpha activity showed a robust positive association with reproduction times (t = 3.085, p = 0.007), with widespread effects across electrodes, indicating that trials with higher-than-average alpha activity were associated with longer reproductions. Theta power similarly showed a positive relationship with reproduction times (t = 3.716, p < 0.001), though this effect was spatially confined to a small set of parietal channels. Regarding aperiodic activity, the offset exhibited a strong positive association with reproduction times (t = 3.578, p = 0.004), while the exponent showed a reliable negative association (t = −3.466, p = 0.002), with both effects primarily localised over frontal electrodes. However, offset and exponent were highly collinear across channels (e.g., Pz: r = 0.91), a potential confounds previously noted in the literature (Donoghue et al., 2020; Ostlund et al., 2022). To evaluate their independent predictive value, we ran two separate LMMs with each aperiodic predictor entered individually. Both parameters positively predicted reproduction times in a small set of posterior channels (offset: P2, P4, PO8, t = 3.383, p < 0.001; exponent: PO8, t = 3.696, p < 0.001), whereas no independent prediction was observed in frontal channels. Given that temporal length has a direct effect on brain–behaviour variability (Buhusi & Meck, 2005; Duffy et al., 2025) we next tested whether the EEG– behaviour association varied as a function of interval duration by incorporating duration as an interaction term (see Methods). No reliable modulation by interval length was found for aperiodic activity (all p > 0.1), indicating that the relationship between EEG activity and single-trial behaviour does not change linearly with interval duration. Only oscillatory alpha activity yielded a few significant frontal electrodes (F1, FC1, Fz, t = 3.441, p = 0.0003). To control for potential time-on-task effects (Benwell et al., 2019; Kopčanová et al., 2025), the trial number was added as covariates in the mixed-effects model (see Methods). Consistent with previous findings (Benwell et al., 2019; Kopčanová et al., 2025), posterior alpha activity increased over the course of the experiment (t = 4.476, p < 0.001; Supp. Fig. 3), while we didn’t find any significant difference in aperiodic activity (p > 0.1). Nevertheless, both oscillatory (alpha: t = 3.012, p = 0.004; theta: t = 3.314, p = 0.001) and aperiodic activity (exponent: t = −3.41, p = 0.003; offset values closely mirrored those of the exponent) remained significant predictors of reproduction times, ruling out the possibility that the EEG–behaviour associations reflect progressive fatigue or neural drift during the experiment (Pessiglione et al., 2025). Finally, given that neural activity varied as a function of interval duration, most notably the decrease in posterior alpha power with increasing interval length, we examined whether the encoding of time was driven by activity at a specific moment within the interval rather than uniformly across it. To this end, the mixed-effects model was repeated using only the first second or only the final second of each encoding interval. We used this approach to dissociate the initial period, during which timing-related effects are not yet expected, from the final period, during which known effects have emerged (e.g., alpha desynchronization for longer intervals; see Fig. 2). When restricted to the first second, alpha maintained a robust positive association with reproduction times (t = 3.183, p = 0.003), with no significant theta effects, and aperiodic activity showed a weaker, spatially limited contribution confined to two frontal channels (offset: CPz, F4; t = 3.38, p < 0.001). In contrast, when restricted to the final second, oscillatory activity was no longer predictive (all p > 0.1), while aperiodic activity showed a significant frontal cluster (exponent: t = −3.437, p = 0.002). This temporal dissociation suggests that alpha and aperiodic activity make distinct contributions at different moments during interval encoding, with alpha dominating early and aperiodic activity dominating late.

**Figure 4:**
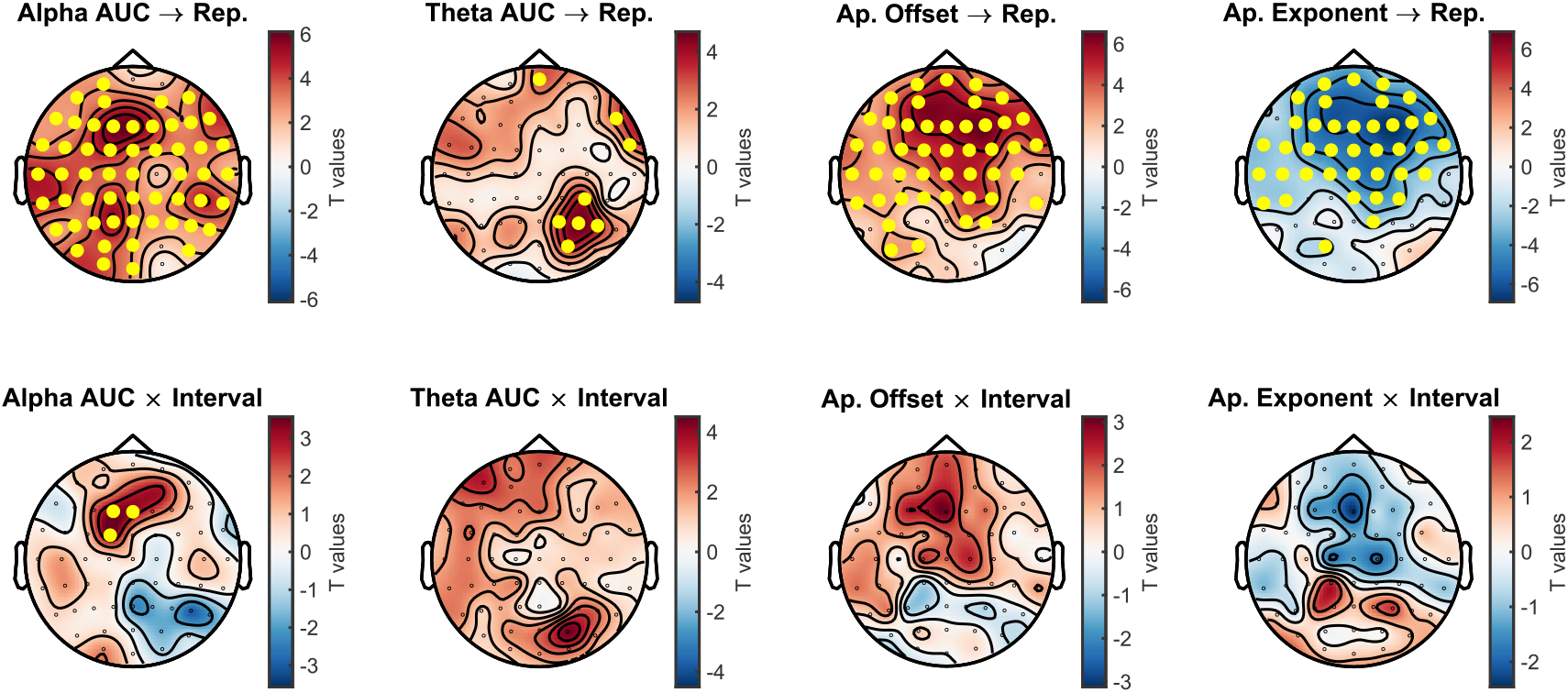
Channel-wise mixed-effects results linking EEG spectral components to reproduction times and their modulation across interval duration. Top row: Scalp topographies of *t*-values from linear mixed-effects models predicting single-trial reproduction times from trial-wise EEG features: Alpha AUC, Theta AUC, aperiodic offset, and aperiodic exponent. Warmer colors indicate positive associations (higher values predicting longer reproductions), whereas cooler colors indicate negative associations. Yellow markers denote channels reaching statistical significance (p < 0.05, FDR corrected). Bottom row: Scalp topographies of *t*-values for the interaction between EEG features and interval duration, indexing how neural– behavior relationships scale with increasing temporal demands

## Discussion

The present study examined whether supra-second timing might be explained by broader neural states that support sustained task engagement. We now turn to discuss three main findings. First, we found that interval encoding was accompanied by a coordinated decrease in posterior alpha power and aperiodic offset. The robust reduction in posterior alpha power during interval encoding is consistent with previous work linking top-down modulation of alpha activity with attentional engagement and temporal expectation (Jensen & Mazaheri, 2010; Nobre & Van Ede, 2018; Sadaghiani & Kleinschmidt, 2016). Critically, this effect co-occurred with a decrease in aperiodic offset, suggesting a coordinated reconfiguration of oscillatory and non-oscillatory processes. Since the aperiodic offset reflects aggregate neuronal firing rates (Miller et al., 2009), its reduction with increasing interval duration points to a progressive suppression of background broadband excitability as the anticipated end of the interval approaches. This interpretation is consistent with evidence that both alpha desynchronization and aperiodic changes occur during the interval preceding an expected event, scaling with attentional anticipation rather than task difficulty per se (Gyurkovics et al., 2022; Kałamała et al., 2024). However, for our knowledge, this is the first time that this aperiodic effect is reported specifically in a timing experiment. Together, the co-occurrence of these two effects may therefore reflect a shared anticipatory reconfiguration of cortical excitability in posterior sensory areas, with both serving as complementary indices of the same underlying neural configuration (Donoghue et al., 2020; Preston et al., 2025). Second, interval duration did not produce robust changes in neural activity during the delay period. Contrary to predictions derived from working memory load accounts (Chikhi et al., 2022; Engen et al., 2026; Frelih et al., 2025; Pavlov & Kotchoubey, 2022; Virtue-Griffiths et al., 2025), we did not observe robust modulation of oscillatory or aperiodic activity during the delay period as a function of interval duration. This absence of interval-dependent modulation suggests that longer intervals do not increase computational demands. That is, interval duration does not contribute to increased cognitive load in the way that, for instance, number of items in working memory do. This finding is also consistent with recent evidence, where alpha power indexes the number of duration items maintained in working memory, rather than the total duration of the interval sequence per se (Herbst et al., 2026). Third, behavioural performance was linked to neural activity at both trait and state levels. At the level of individual differences, we found that higher baseline alpha power robustly predicted steeper behavioural slopes, reflecting more accurate timing. This result aligns with the view that alpha oscillations index tonic alertness and the capacity for sustained cognitive control (Hemmerich, Luna, et al., 2025; Sadaghiani & Kleinschmidt, 2016), and extends it by showing that this capacity relates to the stability of temporal representations. An interesting link to this finding comes from the concept of “readiness to remember”: research on episodic memory has shown that pre-task preparatory attention and goal-state coding (i.e., the neural and cognitive conditions a person brings to a task before it begins) are strong predictors of performance variability both within and across individuals (Madore et al., 2022). Translated to our context, individuals with higher baseline alpha may be better prepared to enter each encoding interval in a stable control state, producing more accurate temporal representations. This is also consistent with evidence that working memory performance depends not only on how much information is held, but on how stably it is maintained in awareness (Allen et al., 2026; Cowan et al., 2024). A further consideration concerns frontal theta activity: growing evidence suggests that clear oscillatory theta peaks are not consistently observable across individuals, complicating the interpretation of frontal midline theta as a stable oscillatory marker of cognitive demand (Engen et al., 2026). Consistent with this view, we found that while oscillatory alpha peaks were present in all but one participant, identifiable theta peaks were observed only in approximately half of the sample. In addition, recent work has suggested that some effects traditionally attributed to frontal theta may instead reflect broader changes in aperiodic neural activity or broadband spectral dynamics (Engen et al., 2026; Herweg et al., 2020; Lu, 2025). For these reasons, we refrain from overinterpreting the null effects of frontal theta: its insensitivity to interval duration during the delay period and its limited contribution to individual differences in timing accuracy. At the trial level, reproduction times were predicted by both oscillatory and aperiodic activity during the encoding interval. First, alpha showed a robust positive association with the reproduced duration, widespread across the scalp. Since interval encoding requires sustained internal monitoring of subjective time rather than external stimulus detection, higher alpha during encoding period may reflect a state of heightened inward attention that amplifies the subjective representation of elapsed duration, producing longer reproductions. Thus, alpha variability accounts for performance both at a trait-level and at a state-level: stable baseline alpha captures individual accuracy in temporal encoding, while trial-level fluctuations reflect momentary variation in the magnitude of the internal time estimate. Examining both levels is therefore essential for a complete characterisation of neural-behavioural relationships (Madore et al., 2020; Rosenberg, 2025; Unsworth et al., 2020; Urai, 2026). With respect to aperiodic activity, the trial-level associations involved both the exponent and the offset, which reflect complementary aspects of the spectral structure: the exponent indexes the steepness of the 1/f fit in log-log space, a measure linked to the ratio of excitatory to inhibitory synaptic currents (Donoghue et al., 2020; Gao et al., 2017), while the offset captures the overall level of broadband neural activity, related to aggregate neuronal firing rates (Miller et al., 2009). The opposing directions of these two effects (positive offset-reproduction and negative exponent-reproduction relationships) might be therefore mechanistically interpretable: on longer-reproduction trials, broadband activity was elevated while the spectral slope was relatively steeper, both consistent with a decreased E/I ratio (Gao et al., 2017). This interpretation aligns with proposals that aperiodic changes index domain-general engagement of control-related networks (Lu, 2025). However, caution is warranted on two counts. The high collinearity between offset and exponent in our data (see Results) limits the ability to draw independent conclusions about their separate cognitive roles. A further finding concerns the temporal structure of neural predictors within the interval. Alpha and aperiodic activity did not contribute equally across the encoding period: alpha power had greater predictive value during the first second of the interval, whereas aperiodic activity emerged as the dominant predictor near the interval endpoint. Rather than contradicting an excitability-based account, this dissociation refines it: early alpha may reflect the onset of a sustained attentional state that sets the general excitability context for temporal encoding, while the late aperiodic shift may capture anticipatory suppression as participants approach the expected interval boundary. Together, both signals contribute to the final reproduction, but at distinct moments and through potentially distinct mechanisms. Whether this temporal structure persists, weakens, or reorganises at substantially longer intervals (e.g., 10 s or more) remains an open question for future research. In conclusion, the dual-level pattern -- the trait differences in baseline and the single trial state fluctuations -- converges on a coherent picture: supra-second timing performance reflects the stability of domain-general neural configurations that regulate sustained conscious engagement, with oscillatory and aperiodic activity serving as complementary windows onto the same underlying excitability state. Furthermore, in line with recent literature (Engen et al., 2026; Frelih et al., 2025; Herweg et al., 2020; Herzog et al., 2024; Preston et al., 2025), the present results suggest that a full account of cognitive timing requires moving beyond a purely oscillatory framework.

## Materials and Methods

### Subjects

A total of 37 healthy individuals (23 women) were recruited for our experiment (mean age = 23.2 ± 2.1 years). Participants were recruited from the community of the Hebrew University of Jerusalem (HUJI) and were compensated for their time with either class credit or monetary reward. All procedures were approved by the Hebrew University of Jerusalem (HUJI) Institutional Review Board (or Ethics Committee) and conducted in accordance with the Declaration of Helsinki. A total of 3 subjects were removed post-hoc due to technical issues in the EEG recording and another 5 subjects were removed for failure to discriminate interval (β < 4SD of the subject distribution). For the final analysis, we used a total of 32 subjects.

### Experimental design

Participants performed a temporal reproduction task. The timing structure of each trial was similar to a previous experiment using supra-second intervals (Kononowicz et al., 2015). At the start of each trial, a red dot in the centre of the screen represented the fixation point, followed by a fixed 1.5s delay. Two consecutive black circles marked the interval to-be-perceived, or Encoding Interval (EI, Fig 1A). Each marker stimulus had a duration of 0.1 s, so the interval to be perceived was the empty gap between them. The interval durations were either 2, 3 or 4 seconds. After another fixed 1.5 second delay, a third stimulus (light-grey circle) determined the onset of the reproduction interval (RI, Fig 1A), where the subject provided the response, and was instructed to reproduce the interval, as accurate as possible. During the gaps between the intervals, the participants fixated a blank black screen. The experiment was composed of 12 blocks, with breaks in between. Each block contained 12 trials, 4 for each interval length. EI durations were randomized within each block. Subjects received a short practice prior to the experiment, using a fixed EI of 1.5 s. Finally, participants were instructed not to use counting strategies during the main experiment.

### EEG acquisition and preprocessing

EEG was recorded using a 62-channel active electrode system (g.Tec, Austria) with electrodes mounted on a standard g.GAMMAcap (g.Tec, Austria) and a g.HIamp amplifier (gTec, Austria). Electrodes across the scalp were distributed according to the international 10–20 system. Signals were acquired using a g.HIamp amplifier and sampled at 512 Hz. Electrode impedances were kept below 5 kΩ. In addition, for all participants we recorded the horizontal electrooculogram (EOG) using passive electrodes placed at the outer canthi of both eyes and the vertical EOG using electrodes placed above and below the left eye. Physiological data and event markers were recorded and synchronized using the Lab Stream Layer protocol (Kothe et al., 2025). Highly noisy F9 and F10 electrodes were removed from the analysis, as previously done. EEG data were preprocessed offline, using Fieldtrip (Oostenveld et al., 2011) and EEGLAB (Delorme & Makeig, 2004) functions. Slow drifts were removed using a custom function previously developed in lab, based on spline interpolation (Ofir & Landau, 2022). This was intended to avoid the potential detrimental effects of filtering, such as the commonly used 0.5 Hz high-pass filter (de Cheveigné & Nelken, 2019). The EEG data were re-referenced to the average of the earlobes. Bad electrodes were removed by visual inspection and reconstructed using ‘spline’ method. The line noises were automatically removed using Zapline-plus algorithm (Klug & Kloosterman, 2022). EEG was low pass filtered at 120 Hz. Data were detrended and demeaned using custom MATLAB scripts. Physiological artifacts (cardiac activity, eye movements, and blinks) were identified through visual inspection and removed using independent component analysis (ICA, mean_comp_rem = 3.780, SD_comp_rem = 1.634). Muscle artifacts were manually removed after visual inspection. Finally, EEG data were transformed with Laplacian filters using the Fieldtrip function *ft_scalpcurrentdensity*.

### Spectral Decomposition and Time-Resolved Parameterization

EEG power spectra were parameterized into periodic (oscillatory) and aperiodic components using the FOOOF algorithm (Donoghue et al., 2020), implemented in Fieldtrip. In all the analysis, PSDs were estimated using single-taper FFT (FieldTrip *mtmfft*, Hanning taper) over a 2–40 Hz frequency range. For the time-resolved analysis, FOOOF was fit within 1 s sliding windows, shifted in 0.1 s steps. FOOOF was applied using a fixed aperiodic mode, with peak width limits set to 1–12 Hz, maximum number of peaks = 3, minimum peak height = 0.10, and peak threshold = 2. Peak detection was restricted to the 3–12 Hz range (theta–alpha band), unless otherwise specified. These parameters were used in previous experiments in attention and working memory (Bender et al., 2025; Engen et al., 2026). For each subject and channel model quality was assessed using the coefficient of determination (R^2^). The quality of fitting in our dataset had a mean R^2^ of 0.789 across participants (10th percentile = 0.729; 90th percentile = 0.846). Oscillatory components were quantified relative to the aperiodic fit. Individual alpha frequency (IAF) was defined as the centre frequency of the largest peak within the 7–13 Hz range. Alpha and theta bands were defined dynamically for each subject and channel: alpha as IAF ± 2 Hz, and theta as 3 Hz to (IAF − 2 Hz). Peaks were considered present only if they exceeded a minimum amplitude threshold. In agreement with previous findings (Engen et al., 2026), we found a clear frontal theta peak in a third of the subjects (Fz: 34.4% of subjects) while posterior alpha was present in all but one participant (31/32). Following previous literature (Bender et al., 2025; Cunningham et al., 2023; Engen et al., 2026) oscillatory power was quantified as the area under the curve (AUC) of the residual spectrum (i.e., the difference between the full model and the aperiodic component in log10 space)

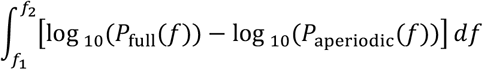

For the FOOOF fitting in sliding window analysis and for the individual trait-analysis, we averaged the PSD across all trials within each channel and within each subject. Following a previous approach (Engen et al., 2026), trials with poor model fits were excluded using a trimming procedure based on the distribution of R^2^ values within each subject, channel, and interval (±2 standard deviations), resulting in the removal of approximately 8% of trials. Analyses run with all trials included yielded similar results. When the ‘knee’ parameter was included in the FOOOF fitting, overall model fit quality decreased; therefore, the fixed aperiodic mode was retained.

### Bayesian model

To characterise individual differences in temporal reproduction performance, we fitted a Bayesian observer model to each participant’s trial-level data. This model decomposes reproduction responses into contributions from a noisy sensory measurement and a prior expectation over interval durations, yielding participant-level estimates of sensory noise (Weber fraction), prior mean, and prior variance. From these estimates, we derived two behavioural summary metrics: a behavioural slope (indexing the gain with which encoded intervals are translated into reproduced durations) and a central tendency index (quantifying the degree to which reproductions are biased toward the prior mean). Prior to fitting the Bayesian observer model, we excluded RT outliers exceeding ±3 standard deviations from the mean per condition. The Bayesian observer model was implemented in Stan using the package brms (Bürkner, 2017). Convergence was confirmed (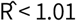 < 1.01; ESS > 400). Individual differences in temporal reproduction were quantified using a subject-wise Bayesian observer model fitted to trial-level data. For each participant, reproduction responses were modelled as arising from noisy sensory measurements (*m* ∼*N*(*t*, (*kt*)^2^)) combined with a Gaussian prior over interval durations (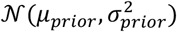). Model parameters (Weber fraction *k*, prior mean μ_*prior*_, prior variance 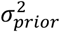) were estimated jointly via Hamiltonian Monte Carlo sampling, with subject-level random intercepts on each parameter. To quantify the difference between individual performance and the veridical reproduction, we computed for each participant a behavioural slope by fitting a linear regression to the mean reproduced durations across intervals (2, 3, and 4 s). This is similar to the regression index, previously described in literature (Cicchini et al., 2012). In parallel, a model-predicted slope was obtained by applying the same regression to the posterior mean estimates derived from the fitted model. Central tendency was quantified based on the relative weighting of sensory evidence and prior information. Specifically, the sensory weight at each interval was defined as 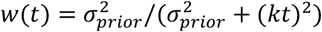, and central tendency was computed as 1 − *w*(*t*). A subject-level central tendency index was obtained by averaging this quantity across intervals, providing a measure of the overall bias toward the prior.

### Statistical analysis

To identify the temporal evolution of oscillatory and aperiodic activity over the course of each trial, we applied a sliding window spectral parameterization approach. EEG power spectra were estimated within overlapping 0.5 s windows (shifted in 0.1 s steps) across the full encoding and delay periods for each participant, channel, and interval condition. Within each window, FOOOF was applied to extract oscillatory (alpha, theta) and aperiodic (offset, exponent) parameters. The resulting time series were then submitted to permutation-based regression analyses to test for systematic modulation by interval duration, separately for the encoding and delay epochs. Within each epoch (final 1 s of the encoding interval; 1.5 s delay period), the metric was averaged across time, and statistical comparisons were performed using cluster-based permutation regression (ft_statfun_depsamplesregrT) implemented in FieldTrip (Maris & Oostenveld, 2007), with 3,000 randomizations and cluster correction controlling for multiple comparisons across channels (minimum 2 neighbouring channels). The sliding-window approach follows Bender et al. (2025). For the trait-level (across-subjects, channel-wise) EEG–behaviour correlations, p-values were corrected using the Benjamini–Hochberg false discovery rate (FDR) procedure (Benjamini & Hochberg, 1995; α = 0.05). Single-trial relationships between aperiodic EEG activity and behavioural responses were assessed using linear mixed-effects models (LMMs) implemented in MATLAB’s fitlme. For each channel, the final reproductions were modelled using the following specification:

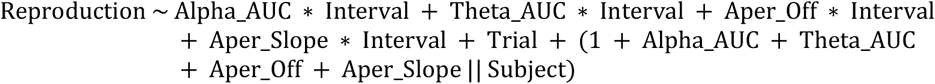

Reproductions and FOOOF parameters were centred within each Subject × Channel × Interval cell. Interval was treated as a continuous linear predictor and mean-centred (re-coded as {−1, 0, +1} for the three encoding durations of 2, 3, and 4 s), so that the main-effect coefficients reflect the average across intervals; each neural predictor was tested in interaction with Interval, and Trial number was included as a fixed-effect covariate to control for time-on-task. Random intercepts and random slopes for all four neural predictors (alpha, theta, aperiodic offset, aperiodic exponent) were included, with an uncorrelated (diagonal) random-effects covariance structure (MATLAB CovariancePattern = ‘Diagonal’), to account for between-subject variability in both baseline reproduction times and sensitivity to neural predictors. For each fixed effect, regression coefficients, t-statistics, and p-values were extracted and FDR-corrected across channels. In all the statistical reports of the LMM (t-values, betas and p-values), the values are the average of the parameters across the significant channels.

## Supporting information

Supplemenary_TPMms

## Acknowledgement

We thank Yoel Gordon for the contribution on the first part of the experiment development and the data acquisition. This project has received funding from the European Research Council (ERC) under the European Union’s Horizon 2020 research and innovation programme (grant agreement no. 852387).

