## Supplementary material for "Supra second timing reflects oscillatory and aperiodic EEG dynamics": Supplemenary_TPMms

### Supplementary materials

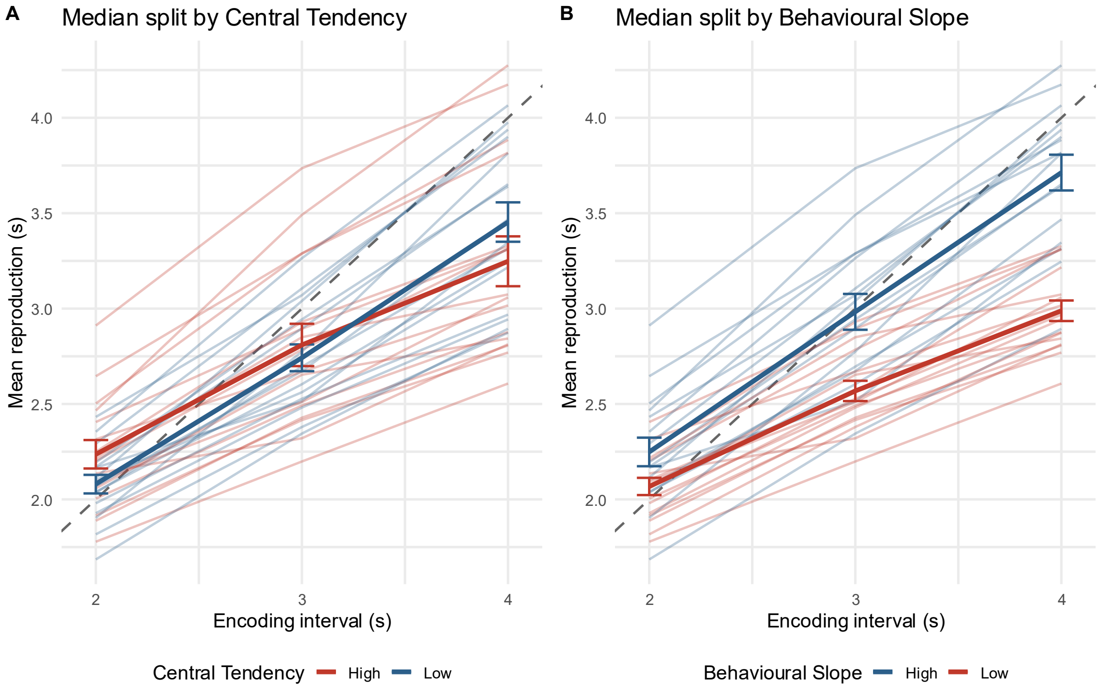


**Supp. Fig. 1: behavioral reproduction performance stratified by central tendency and predicted slope.**

Mean reproduction times as a function of encoding interval (2, 3, 4 s) shown separately for two median-split groups. Left: Participants split by central tendency (CT). High-CT individuals (red) exhibit a stronger bias toward the mean interval, reflected in a shallower slope, whereas low-CT individuals (blue) show more veridical scaling with interval duration. Right: Participants split by predicted slope from the Bayesian observer model. High-slope individuals (blue) display more accurate interval reproduction (steeper scaling), while low-slope individuals (red) show greater compression toward the mean. Thin lines represent individual subjects, thick lines indicate group means, and error bars denote ± 1 SEM. The dashed unity line indicates veridical performance.

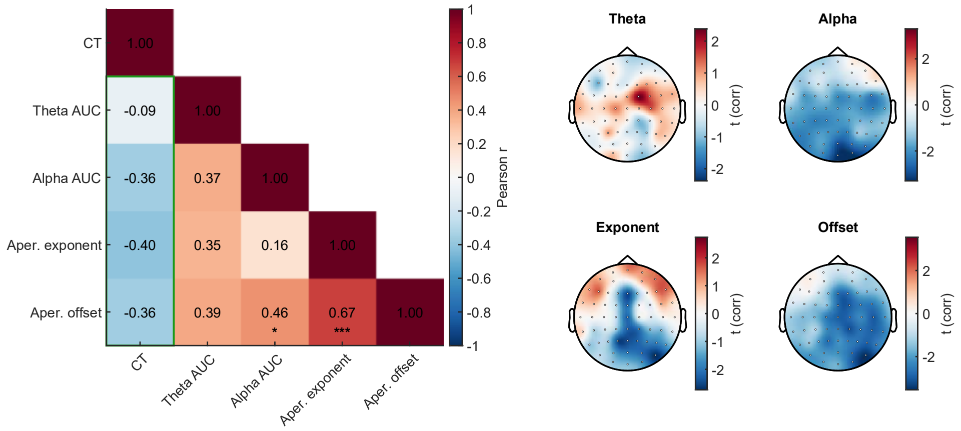


**Supp. Fig. 2: Relationship between central tendency and EEG spectral components.**

Left panel: Pearson correlation matrix across subjects (N = 32) including central tendency (CT), theta power (Theta AUC), alpha power (Alpha AUC), aperiodic exponent, and aperiodic offset, averaged across a posterior electrode cluster (CPz, P1, Pz, P2, POz). Green-outlined correlations are represented in the topographies on the right. Significant correlations are denoted by asterisks (* $p < 0.05$, ** $p < 0.01$, *** $p < 0.001$; FDR corrected). Right panels: Scalp topographies of channel-wise correlation t-values between central tendency (CT) and EEG metrics. Top row: correlations with Theta AUC (left) and Alpha AUC (right). Bottom row: correlations with aperiodic exponent (left) and aperiodic offset (right). Yellow markers denote channels reaching statistical significance ($p < 0.05$, FDR corrected).

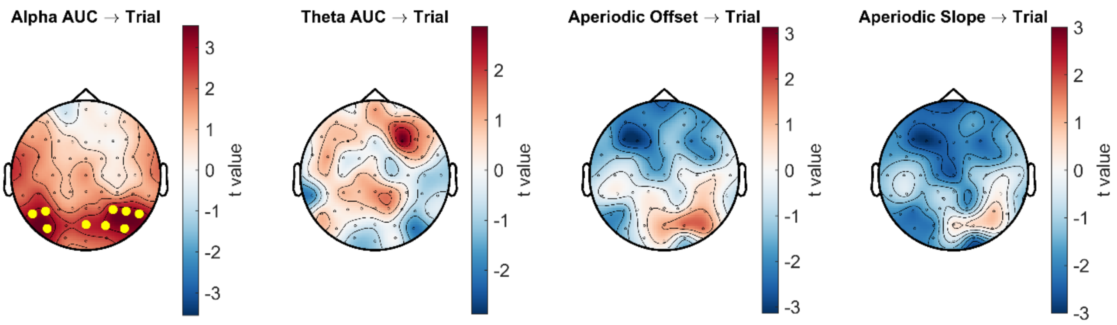


**Supp. Fig. 3: trial-level effects of EEG spectral components and their relationship with reproduction times.**

Top row**:** Scalp topographies of *t*-values from linear mixed-effects models assessing the effect of trial progression (trial index) on EEG features: Alpha AUC, Theta AUC, aperiodic offset, and aperiodic exponent (slope). Warmer colors indicate increases across trials, whereas cooler colors indicate decreases. Yellow markers denote significant channels ($p < 0.05$, FDR-corrected). Summary statistics (right) indicate the number of significant channels for each feature, revealing prominent trial effects for aperiodic components, particularly the exponent. The Trial → RT map reveals a global positive effect, indicating increasing reproduction times over the course of the experiment.
